# Niche partitioning as a selective pressure for the evolution of the *Drosophila* nervous system

**DOI:** 10.1101/690529

**Authors:** Ian W. Keesey, Veit Grabe, Markus Knaden, Bill S. Hansson

**Affiliations:** Max Planck Institute for Chemical Ecology, Department of Evolutionary Neuroethology, Hans-Knöll-Straße 8, D-07745 Jena, Germany

## Abstract

The ecological and developmental selective pressures associated with evolution have shaped most animal traits, such as behavior, morphology and neurobiology. As such, the examination of phylogenetic characteristics of the nervous system can be utilized as a means to assess these traits, and thus, to evaluate the underlying selective pressures that produce evolutionary variation between species. Recent studies across a multitude of *Drosophila* have hypothesized the existence of a fundamental tradeoff between two primary sensory organs, the eye and the antenna. However, the identification of any potential ecological mechanisms for this observed tradeoff have not been firmly established. Our current study examines two monophyletic species within the *obscura* group, and asserts that despite their close relatedness and overlapping ecology, they deviate strongly in both visual and olfactory investment. Here we contend that both courtship and microhabitat preferences mirror and support the observed inverse variation in these sensory traits. Moreover, that this variation in visual and olfactory investment between closely-related species seems to provide relaxed competition, a process by which similar species can use a shared environment differently and in ways that help them both coexist. The nervous system has a unique role in evolution as it provides the functional connection between morphology, physiology, and behavior. As such, characterizing any tradeoffs between costs and benefits for the nervous system may be essential to our understanding of animal diversity, as well as vital to our understanding of the selective forces that have shaped the natural world.

## INTRODUCTION

The genus *Drosophila* provides an incredible array of phenotypic, evolutionary and ecological diversity [1–5]. Several thousand relatives of the model organism *D. melanogaster* inhabit all continents except Antarctica, and occur in almost every type of environment. Due to their vast variation in behavioral, morphological and natural history traits, the comparison of Drosophilid flies provides an enormous potential for the understanding of driving forces in evolutionary processes. In particular, the feeding, courtship and breeding sites of this genus are tremendously diverse, including both generalists and specialists, and spanning extreme dietary variation and host utilization such as different stages of fruit decay, as well as flowers, mushrooms, sap or slime flux, rotting leaves, cacti and many other sources of microbial fermentation. It is important to note that preferences in feeding and oviposition have shifted numerous times, and closely related species are known to utilize different types of food resources [6,7], or to visit a host at different stages of decay [8–10]. At the same time, it is common to find phylogenetically distant species using the same host, or living in overlapping environments [11–15]. Therefore, the spatial distribution of species over discrete patches of an ecosystem, such as within a temperate forest, might vary according to discrete microhabitats. Moreover, these local environments may create different selective and energetic pressures for neuroanatomy [16] that in turn could provide an opportunity and rationale for the coexistence of many species within a single habitat or ecological niche [17,18].

In an earlier paper we showed that robust idiosyncrasies exist between visual and olfactory investment across this genus [2], including many examples of inverse variation within a subgroup, and between sympatric species that utilize seemingly identical host plants or food resources. However, most species have an understudied ecology, and other than information about where and when they were collected for laboratory establishment, we often know very little about their natural habitats or ecological preferences. The aim of the present paper is to determine whether behavioral, phenotypic, and neuronal differences between close relatives all combine to support the coexistence of different species within a single ecological habitat. More specifically, we document that these sensory traits vary significantly between two close relatives within the *obscura* group – *D. subobscura* and *D. pseudoobscura* – and we examine in detail the potential driving forces of speciation, such as biotic and abiotic factors. We assert that even between close, phylogenetic relatives, these differences in sensory investment are strongly apparent, and we propose that these differences could reduce interspecies competition via resource partitioning and through innate variation in microhabitat preferences, thus promoting speciation and stabilizing selection using natural sensory trait variation.

## RESULTS

### External morphology of sensory systems

In order to examine the sensory traits of the two closely related and sometimes co-occurring species – *D. subobscura* and *D. pseudoobscura* – we quantified their visual and olfactory investment by first measuring the external morphology of their visual and olfactory systems. Eight to ten females of the two species were photographed using a Zeiss AXIO microscope, including lateral, dorsal, and frontal views. We then measured across a variety of physical characteristics, such as surface areas of the compound eye, antenna, maxillary palps, ocelli, and overall body size, as well as head, thorax, abdomen and femur length. We also generated metrics for the number of ommatidia as well as measures of trichoid sensilla for each species. In general, we found that *D. subobscura* possessed much larger eyes in regards to surface area, as well as 25-30% more ommatidia than its close relative, *D. pseudoobscura*, though ommatidia diameter was identical (**Figure 1A-E; Supplementary Figure 2**). While there was some variation in individual size within and between species (with *D. subobscura* exhibiting larger dimensions in all measured body parts; **Supplementary Figure 1A-C**), we note that eye surface area was consistently correlated with ommatidium number **(Figure 1C)**, suggesting that eye surface area provides a good approximation of visual investment. Here we note that both species had a nearly identical linear relationship between surface area and number of ommatidia (**Figure 1C**), with *D. subobscura* possessing larger eyes. While *D. pseudoobscura* possessed smaller eyes and a reduced ommatidum count, this species instead displayed larger antennal surface areas relative to *D. subobscura* females (**Figure 1A,F**). Interestingly, not all metrics related to sensory organs on the head were different between these closely related species. For example, the maxillary palps (**Figure 1G**) did not display any significant variation in surface area between the species, but we do note differences in the ocelli (**Supplementary Figure 2**). Thus, these changes to sensory systems on the head appear mostly restricted to the antenna and to the visual modalities.

**Figure 1.**
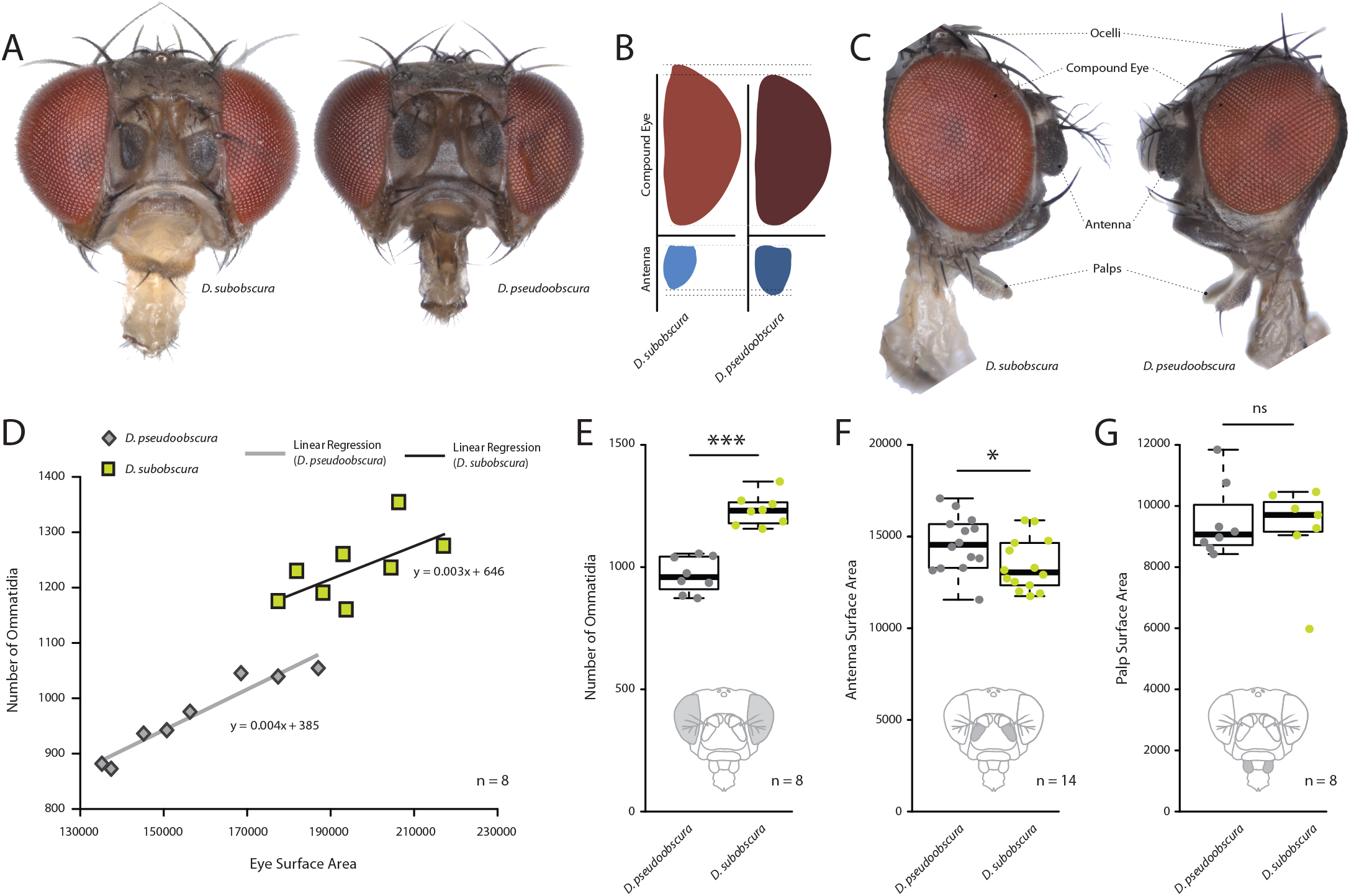
Comparative morphology of external sensory systems. (A) Examples of frontal head replicates, where eye and antenna surface area was measured. Note the differences in pigmentation, as well as the size of the compound eye and third antennal segment from both species. (B) Side-by-side comparison of both the compound eye (red) and third antennal segment (blue) from each fly. (C) Example of lateral views used for measurements, including compound eye surface area, antennal surface area, and maxillary palp surface area. (D) Intra- and interspecies correlations between eye surface area and the number of ommatidia for *D. subobscura* (yellow) and *D. pseudoobscura* (grey). (E-G) Comparison of ommatidia counts (E), antennal surface (F), and palp surface (G) for both species. Boxplots represent the median (bold black line), quartiles (boxes), as well as the confidence intervals (whiskers). Mann-whitney U test; ***, p<0.001; *, p<0.05; ns, p>0.05.

### Comparative neuroanatomy of visual and olfactory investment

As we had already established divergent external morphology between the species, especially in regard to vision and olfaction, we next focused our attention on the primary processing centers in the brain for these sensory systems, including the antennal lobe (AL) and optic lobe (OL) (**Figure 2AB**). After correcting for adult size (using the hemisphere or central brain volume as a reference for each species), we identified a relative increase of the AL size for *D. pseudoobscura* (**Figure 2C**), as well as a relative decrease of the size of its OL (**Figure 2D**) when compared to the same neuropils for *D. subobscura* adults. These inverse values between the two sensory systems correspond strongly to the variations we measured in the external morphology, where one species had larger eyes but smaller antenna, and vice versa. Moreover, to highlight the regions of the OL that show the largest increases, we provide similar metrics for relative size for the lobula plate, lobula and the medulla (**Figure 2E**), where all brain regions (again when corrected for total brain size) are bigger in *D. subobscura*, but only the medulla is significantly larger.

**Figure 2.**
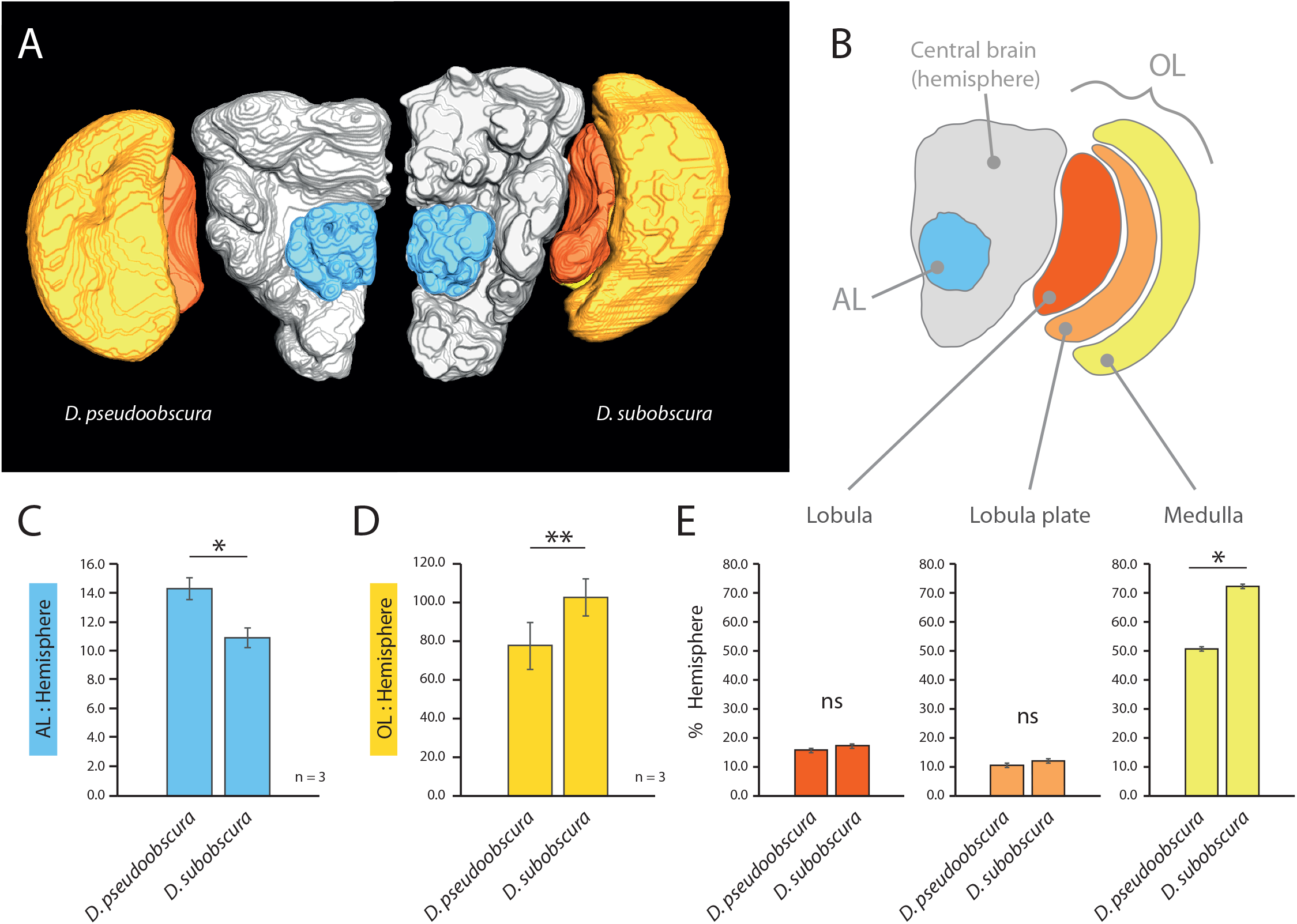
Comparative morphology of primary processing centers in the brain. (A) Three-dimensional reconstructions of the neuropils of *D. pseudoobscura* and *D. subobscura* adult females. (B) Diagrammatic representation of the brain, with color-coded and labeled volumetric sources. Antennal lobe, AL, blue; hemisphere, grey; optic lobe, OL, with medulla (yellow), lobula plate (orange) and lobula (red). (C-E) Relative size of AL (C), OL (D), and lobula, lobula plate, and medulla (E) as compared to respective hemisphere [%]. Boxplots represent the median (bold black line), quartiles (boxes), as well as the confidence intervals (whiskers). Mann-Whitney U test; **, p<0.01; *, p<0.05; ns, p>0.05.

### Courtship and mating behavior differences between *obscura* species

In order to ascertain the possible ramifications of inverse eye and antenna variation between our two species, we proceeded to examine behaviors related to mate selection and courtship. Previous research has shown that *D. subobscura* displays light-dependent courtship, and will not successfully copulate in the dark [19–21]. Counter to this, *D. pseudoobscura* mating is light-independent, and courtship can successfully occur regardless of light conditions [20–24]. Therefore, as we wanted to observe and dissect the behavioral motifs and succession of events that lead to successful courtship, we performed courtship trials under identical conditions for both species. We recorded videos using virgin males and females that were introduced into a small courtship arena (**Figure 3**). Several differences were immediately noted between the species. *D. pseudoobscura* males oriented themselves either behind or to the side of the female during courtship, often forming a right angle to her with the male head focusing on the last few abdominal segments of their potential mate (**Figure 3A**). Next, this species performed characteristic wing vibrations and singing, with the outstretched wing always nearest to and in the direction of the head of the female (**Figure 3A**), and with the male constantly in pursuit of the female from behind or from the side. In stark contrast, observations of the courtship of *D. subobscura* showed that the males of this species often dart around in a circular arc to put themselves directly in front of the path of the female, and appear to arrest her movement (**Figure 3B,C**). This frontal positioning by the *D. subobscura* male results in most of the subsequent courtship behaviors occurring in front of the female and within her visual field, including the male wing displays. Here, *D. subobscura* was not observed to vibrate the outstretched wing (unlike *D. pseudoobscura* males, which are known to sing), and instead, seems to angle or tilt the outstretched wing during the display, possibly as a flash of color or visual display for the female (**Figure 3B,D**).

**Figure 3.**
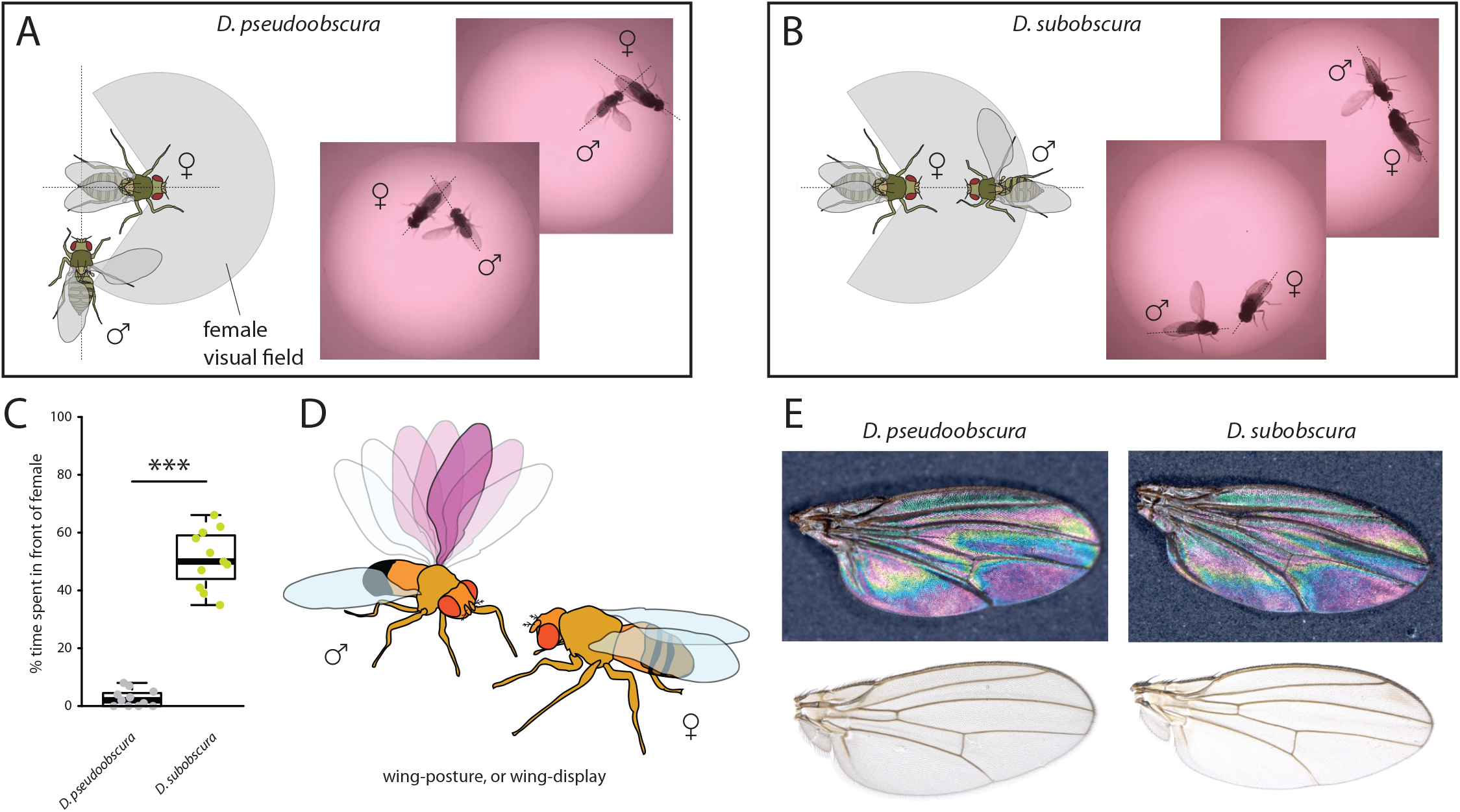
Courtship and mating behavior. A-B) Images of courtship for *D. pseudoobscura* (A) and *D. subobscura* (B) with a schematic of the behavior, highlighting whether the male is in- or outside of the female’s predicted visual field. (C) Time that males spent within the female visual field during courtship. Boxplots represent the median (bold black line), quartiles (boxes), as well as the confidence intervals (whiskers). Mann-Whitney U test; ***, p<0.001 (D) Diagram of *D. subobscura* wing display by the male, where no wing vibration was observed, and instead, a discrete range of wing angles was presented and maintained towards the female mating partner during courtship. (E) Stable structural wing interference patterns observed across the otherwise clear wings of males of both species.

### Phototactic responses by close-relatives of the *obscura* group

Given that we had established that differences in compound eye and antenna sizes correlated with differences in courtship behavior, we next examined whether the morphological investments played any additional role in ecological decisions related to environmental preferences. Here we utilized a simple Y-tube two-choice behavioral assay, where adult flies from each species could select between a light or dark environment (**Figure 4A**). We observed that the smaller-eyed *D. pseudoobscura* significantly preferred to enter the Y-tube arm that was in shadow and darkened (**Figure 4B**). In contrast, the larger-eyed adult *D. subobscura* significantly preferred the Y-tube arm that was in full light.

**Figure 4.**
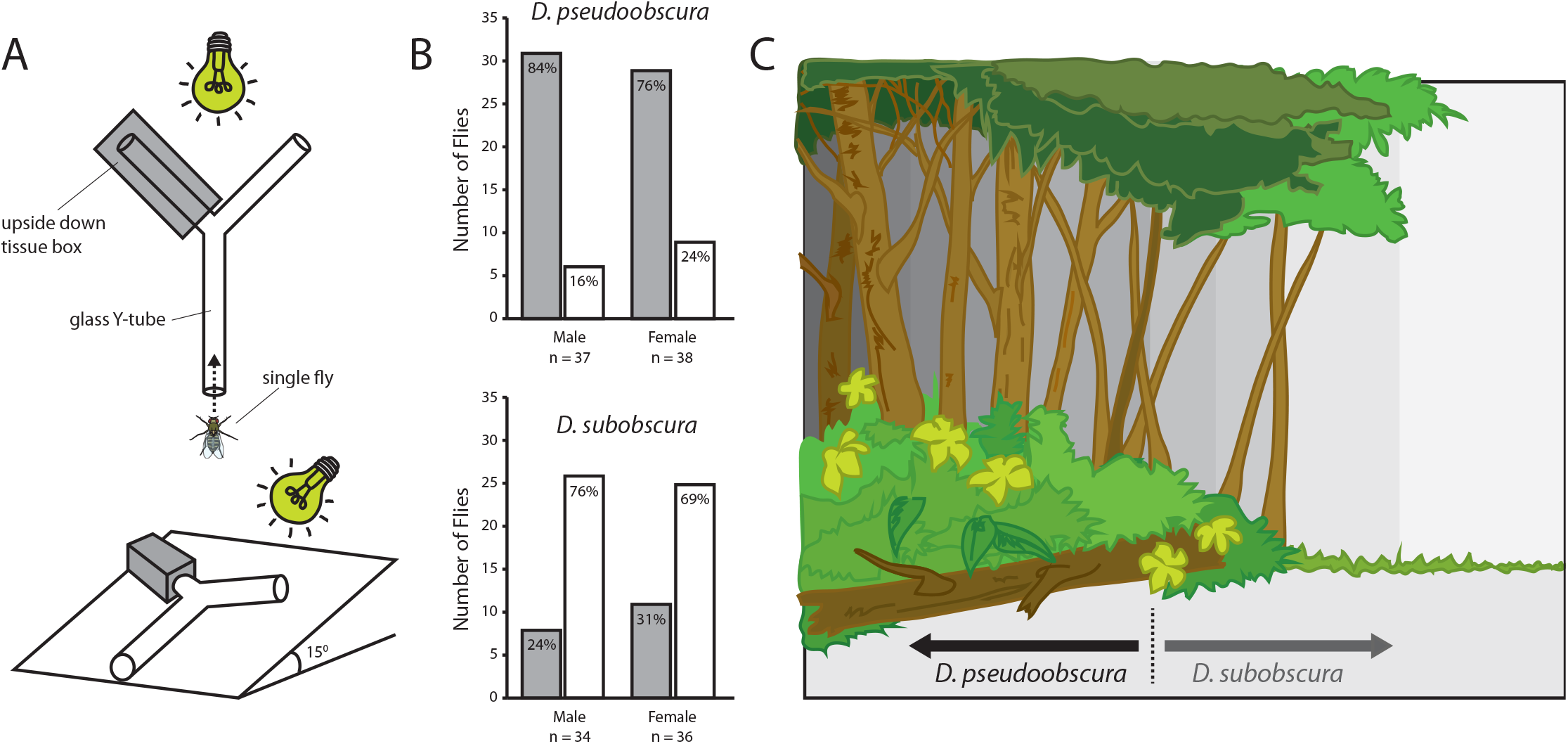
Light preferences and hypothesized niche partitioning by both species. (A) Two diagram views of the Y-tube phototactic response paradigm. Single flies were allowed to choose between either a well-lit or a darkened arm of a Y-tube. (B) Percentages of male and female flies of both species choosing the well-lit or darkened arm of the y-tube. (C) Diagram of ecological niche partitioning where our closely-related *Drosophila* species divide spatially across microhabitats within the same environment, and where light gradients act as an isolation barrier. Here we propose that these two species, despite sharing a forest ecology, create a reduction in either host resource or mating competition via their different preferences towards edge and open canopy environmental conditions, as related directly to their innate preferences for light intensity.

## DISCUSSION

When we imagine examples of isolation barriers, we often consider those that are distinctly physical in nature, such as a mountain range or a remote island biogeography. However, sensory isolation barriers also exist, including differences in pheromone chemistry between geographically overlapping species, or variations in the songs and auditory repertoires of crickets, frogs and birds. In this study, we hypothesize that sources of light gradients may also create strong selective pressures and isolation mechanisms that in turn lead to speciation events or stabilizing selection for opposing phototaxis within otherwise overlapping habitats, such as arboreal forests. These forest microhabitats have been addressed previously as sources for spatial separation between species, including studies directly related to the field-sampling of members of the family Drosophilidae [13,25,26], often with the division of species occurring in proximity to the forest edge. While the evolutionary selective pressures and their effects on the relative size of various components of the nervous system have not been previously examined, it has been suggested that sources of light may be one of the ambient forces driving the observed tradeoff in the evolution of these two sensory structures [2].

Here we demonstrate that two monophyletic species within the *obscura* group, despite being close relatives, deviate strongly in regards to both eye and antenna morphology (**Figure 1**), as well as in their corresponding neuropils for vision and olfaction (**Figure 2**). In addition, we observe that this variation in sensory systems positively correlates with both courtship behavior (**Figure 3**) and environmental habitat preferences (**Figure 4**), especially as related to the relative importance of visual stimuli or sources of light, which appears to be of opposing value between these two sibling species. Previous work has documented this tradeoff or inverse resource allocation between vision and olfaction across more than 60 species within the *Drosophila* genus [2], but the ecological mechanisms and selective pressures underlying this divergence have not been studied explicitly in monophyletic species. While little ecological information is available for a majority of the non-*melanogaster* species, it has been shown repeatedly that many of the members of the *obscura* species group overlap geographically as well as ecologically in their utilization of temperate forest ecosystems [27–29]. In addition, there has been documentation of a geographical overlap between *D.pseudoobscura* and *D.subobscura* in North America, which might make for an ideal field study in the future to test our hypotheses regarding environmental partitioning [37,40]. It would also be important to address additional *Drosophila* species within the *obscura* group, such as *D. affinis, D. persimilis, D. obscura, D. ambigua* and *D. helvetica*, especially in cases where these species potentially directly share ecological overlap in habitat utilization, and where genomic information is perhaps readily available for additional analyses. Here, we test the hypothesis that close insect relatives may divide host or habitat resources through niche partitioning by inversely prioritizing the relative importance of visual stimuli as compared to those that are olfactory. This explanation would be consistent with previous observations that sympatric species often possess inversely correlated eye and antenna sizes despite being close relatives and despite sharing seemingly identical hosts and environmental preferences [2].

Microhabitats often arise in nature, as landscapes are inherently non-uniform [11,30]. These ecological subdivisions have been examined in regards to cline variation or altitude [27,31,32], as well as temperature gradients or differences in water availability [30,32–34]. In addition, several studies have addressed microhabitat variation and its effects on species richness or biodiversity. Moreover, that plant hosts and other nutritional resources such as fungi and yeasts can differ greatly between forest edge and forest interior [26,29,33,35]. Thus, it is well recognized that flora and fauna can vary in both their relative abundance as well as their innate preferences across microclimates within a single habitat, where the fitness of a species is intimately tied to its ability to compete for resources within its own environmental niche. However, the mechanisms by which these innate animal preferences for microhabitats can generate evolutionary pressures or speciation events has not been as thoroughly documented, least of all in a model organism where robust molecular genetic toolkits are available, such as those afforded by the *Drosophila* genus.

Other insects have been examined for their differences in circadian rhythm or phenology, either in regards to host search or mating behaviors, where close relatives are able to reduce competition by varying their activity cycles [36]. Conversely, it has been well-documented that both *D. pseudoobscura* and *D. subobscura* share similar crepuscular activity [25,27,29,37], thus at this juncture we do not feel that the circadian rhythms or seasonal activities of feeding or courtship play any distinct role in the observed evolutionary divergence in relative eye or antenna size. As such, while the light-dependent courtship of *D. subobscura* suggests a difference in daily patterns of mating, this has not been shown [25,27,37]. Thus, we suggest that it is more likely that *D. subobscura* simply uses visual stimuli as a species-defining trait, perhaps initiated via a preference for a better-lit arena to perform their courtship ritual and to attract a potential mate. This would include such microhabitats as a forest edge or an open forest canopy (**Figure 4C**), where visual elements of courtship such as wing displays would be more optimally employed for species identification and female sexual selection given the increases in light availability. Here, we suggest that while both species are assumed to be linked via a forest ecology [27–29], that *D. pseudoobscura* may be more likely to prefer darker, inner-forest habitats, while *D. subobscura* would prefer the forest edge or sections of open canopies within the same forest environment (**Figure 4C**). This light preference would therefore create opposing spatial regions of highest abundance, where each species would reduce overlap with the other by tuning their sensory systems towards either larger-eyes and positive phototaxis, or smaller eyes and negative phototaxis. Moreover, that this shift in the nervous system would then affect both courtship and host preference.

The utilization of forest openings are well studied in avian biology, where males often construct and clear elaborate arenas to perform intricate visual displays for females (i.e. the genus *Parotia* or six-plumed birds of paradise) [38]. However, to our knowledge, the visual capabilities across vertebrate animal species has never been compared to examine evolutionary investments in the nervous system that correlate with visual courtship. Again, we feel it is likely that investment in the visual system might mirror the tendency of a *Drosophila* species to possess a positive phototaxis, which we show between these two *obscura* species, though it remains unclear which of these behavioral phenotypes occurs first and subsequently drives a correlation in the other trait over the course of evolutionary time. It is also unclear which are the most important factors in the visual displays of *D. subobscura* during courtship, for example, whether outstretched wings provide a specific color or UV pattern, or whether this wing display simply generates a flash of bright light reflected towards the female. Moreover, it has been shown that *D. subobscura* does not sing, and thus do not vibrate their wings during display, but we do observe midleg tapping or drumming, which may instead be the auditory component of their courtship ethology. Thus far, no research has simultaneously compared visual and auditory neurobiology or development for these species, but future work should attempt to encompass further sensory modalities. Additional studies will also need to address which photoreceptors are expanded in the compound eye of *D. subobscura* and how they validate the increases in ommatidia when compared to other close relatives. However, previous research has already shown an expansion of the fruitless positive cells in the optic lobes of *D. subobscura* as compared to *D. melanogaster* [39]. Thus, while this pathway has not been addressed yet in *D. pseudoobscura* or any other members of the *obscura* group, it is perhaps again indicative of an evolutionary investment in visual modalities for courtship success, given the visual connection to this fruitless labeled pathway. Nevertheless, it is apparent from our current data that variation in visual and olfactory sensory system development occurs for more than just mating purposes, and appears to match ecological deviations in behavioral phototaxis and microhabitat preferences within a shared ecological niche. In the future, it will continue to be important to test our theories related to niche partitioning as an evolutionary force for sensory variation across other *Drosophila* groups beyond *obscura* and to continue to provide ecological explanations for the observed variation or tradeoff between these two sensory systems in relation to geographical overlap as well as across more species throughout the entire genus.

## Supporting information

Supplementary Video 1 courtship (D. subobscura)

Supplementary Video 2 courtship (D. pseudoobscura)

## ACKNOWLEDGMENTS

This research was maintained through funding provided by the German government and the Max Planck Society (Max Planck Gesellschaft). Wild-type flies were obtained from the San Diego Drosophila Species Stock Center (now The National Drosophila Species Stock Center, Cornell University). We express our gratitude to S. Trautheim and D. Veit for their technical support, expertise and guidance at MPI-CE. We also thank R. Stieber for her expertise and guidance in regards to immuno staining for these two novel fly species.A special thankyou as well to J. Balma for her expert assistance and proficiency in compiling the curated courtship video examples.

## AUTHOR CONTRIBUTIONS

I.W.K. generated the original hypotheses for this study, with support from V.G., B.S.H. and M.K. Photos and measurements of external morphology were conducted by I.W.K. Neuropil dissection and staining was performed in association with V.G. and I.W.K., as were the measurements and reconstructions of primary olfactory and visual organs. Courtship videos and other behavioral data were collected and analyzed by I.W.K. All diagrams, illustrations and figures were created by I.W.K., as was the original written manuscript. Each author subsequently improved the final version of the written document and figures, and assisted with any revisions towards the final publication.

## DECLARATION OF INTERESTS

The authors declare no competing interests.

## STAR ★ METHODS

### METHOD DETAILS

#### External morphometrics from head and body

For each fly species, 8–10 females were photographed using a Zeiss AXIO Zoom.V16 microscope (ZEISS, Germany, Oberkochen), including lateral, dorsal, and frontal views. Flies of each wild type were dispatched using pure ethyl acetate (MERCK, Germany, Darmstadt). Lateral body (40×), dissected frontal head (128×), and dissected antenna views (180×) were acquired as focal stacks with a 0.5x PlanApo Z objective (ZEISS, Germany, Oberkochen). The resulting stacks were compiled to extended focus images in Helicon Focus 6 (Helicon Soft, Dominica) using the pyramid method. Based on the extended focus images, we measured head, thoracic, abdominal, foreleg (femur), as well as funiculus and compound eye surface areas, where all measurements are in µm (Figure 1; Supplementary Figure 1). We also measured surface areas of the maxillary palps and length of the ocelli from both species; however, we did not find any significant difference for the palps (Figure 1C,G; Supplementary Figure 2). Measurements of all body regions were conducted manually using the tools available in Image J (Fiji) software. All raw data available with online version of the manuscript.

#### Ommatidia counts and compound eye surface area metrics

In order to count ommatidia, the compound eye of each species was arranged laterally and perpendicular to the AXIO Zoom.V16 microscope. A total of 8-12 individuals per species were utilized, with only the best 8 specimens used where the eye was completely intact and in focus, where counts were done manually using Image J (Fiji) software tools (Figure 1C-E). We also examined the association between eye surface area and ommatidia counts (Figure 1D). Here we note that both species share nearly identical linear regressions analyses between the number of ommatidia and the associated surface area, thus we conclude that ommatidium diameter is identical between the two species. Though we observed small variations in absolute body size within our species populations that appeared to be correlated with rearing density (e.g. high density produced smaller flies), we also observed a consistently conserved ratio between the eye and antenna morphology regardless of adult body size (data not shown). However, to control for density-dependent plasticity, we maintained both species at a consistent population size (15 females per rearing vial).

#### 3D reconstructions and neuropil measurements

In order to assess neuroanatomy, the dissection of fly brains was carried out according to established protocols [2]. The confocal scans were obtained using confocal laser scanning microscopy (Zeiss confocal laser scanning microscope [cLSM] 880; Carl Zeiss) using a 40x water immersion objective (W Plan-Apochromat 40×/1.0 DIC M27; Carl Zeiss) in combination with the internal Helium-Neon 543 (Carl Zeiss) laser line. Reconstruction of whole OLs and ALs was done using the segmentation software AMIRA version 5.5.0 (FEI Visualization Sciences Group). We analyzed scans of at least three specimens for each and then reconstructed the neuropils using the segmentation software AMIRA 5.5.0 (FEI Visualization Sciences Group). Using information on the voxel size from the cLSM scans as well as the number of voxels labeled for each neuropil in AMIRA, we calculated the volume of the whole AL as well as the individual sections of the OL and the central brain (where central brain values exclude the AL volume).

#### Analyses of courtship and mating behavior

For the analysis of courtship behavior, the adult flies were collected as pupae into single vials (using a wet paint brush), and then later identified by sex after subsequent eclosion. Adults were kept in these single vials for 2 – 6 days after eclosion with access to food and water. Temperature controlled chambers were used for courtship conditions. Here we optimized the temperature for both *obscura* species, where courtship initiation and success was observed to be highest between 18-24 degrees Celsius, which was a substantially lower temperature than previous examinations of *D. melanogaster* courtship. In the behavioral assays, a female fly was first aspirated into the tiny chamber, and secured with a clear cover slide (Supplementary Figure 2G). Next, a male fly was introduced into the same chamber, and video recording was initiated. The flies were recorded under white light illumination for 10 – 15 minutes. If no initiation of courtship was observed after 10 minutes, then videos were halted and new flies were introduced as a novel pair. Videos of successful courtship and copulation were analyzed with BORIS (http://www.boris.unito.it/).

#### Wing interference patterns and pigmentation

In order to assess visual elements of adult wings from both *obscura* species, individual wings from each species were photographed using an AXIO microscope, as was described previously for external head and body metrics. Both clear as well as dark, opaque backgrounds were used to examine wing interference patterns (WIPs) and any other elements of visual information that the wings represent during courtship display (Figure 3E). Here we noted differences in wing shape, as well as sensillum and hair lengths along the wing margins of these two species. However, we did not observe any obvious differences in WIP, nor did we note any apparent differences in pigmentation, color or other visual structures. Thus, it would appear that the wings of the two species are nearly identical, and that perhaps only the behavioral utilization of the wing differs between these species during male courtship (Figure 1A-D).

#### Phototaxis behavior and Y-tube two-choice experiments

A glass Y-tube was fixed and positioned at approximately a 15-degree slope (which encouraged upward walking), with one arm covered with an opaque cardboard box that was cut to match the diameter of the glass (Figure 4A). This cover area provided a heavily darkened arm of the Y-tube, while the other arm was fully illuminated. Both terminal ends of the Y-tube contained sealed glass containers for insect collection and removal. Adults were introduced into the base of the glass Y-tube using an aspirator, where adults could freely walk out of the aspirator pipette tip once they had calmed, and acclimatized to the setup (this greatly reduced escape responses, and random choices). We positioned a light source that mimics natural sunlight wavelengths at the end of the Y-tube, and all overhead illumination (as well as all other sources of light in the chamber) were eliminated. Adult flies were allowed to walk up the Y-tube where they had to then choose between either a dark or light arm, where the first choice was noted for each individual fly (Figure 4B), and time duration was also recorded (Supplementary Figure 1D). After every 10 individuals, an additional, clean glass Y-tube was used (to avoid any contamination from cuticular hydrocarbons or frass/feces left behind by previous flies [41]), and the Y-tubes were rotated after every fly to eliminate any directional bias that could be caused by imperfections in the glass or Y-tube arms. We also rotated the darkened arm every time we exchanged the Y-tube for a clean one, to eliminate any left-right bias. Each day we cleaned glassware with hot soapy water, then rinsed with cold water, then rinsed with ethanol, and lastly we heated them for several hours at 200 degrees Celsius before use in these behavioral assays. In both species, the males showed a stronger trend of light preference than females; however, this trend was not significant (**Figure 4B**). We also noted no significant differences in the time it took flies to make a choice (**Supplementary Figure 1D**), but there was a trend that *D. pseudoobscura* were slighty faster, as were the males of both species when compared to females.

#### Statistical assessments and figure generation

All images and drawings are originals, and were prepared by the first author for this publication. Figures were prepared via a combination of R Studio, Microsoft Excel, IrfanView v4.52, ScreenToGif, VLC Media Player, and Adobe Illustrator CS5. Statistics were performed using GraphPad InStat version 3.10 at α = 0.05 (*), α = 0.01 (**), and α = 0.001 (***) levels. Error bars for bar graphs are standard deviation. Boxplots represent the median (bold black line), quartiles (boxes), as well as the confidence intervals (whiskers).

#### Supplementary Information

All data supporting the findings of this study, including methodology, display examples, raw images and z-stack scans, statistical assessments, courtship videos, as well as other supplementary materials are all available with the online version of the manuscript.

**Supplementary Movie 1.** Courtship behavioral video clip examples for *D. subobscura* adults.

**Supplementary Movie 2.** Courtship behavioral video clip examples for *D. pseudoobscura* adults.

---

**Supplementary Figure 1.**
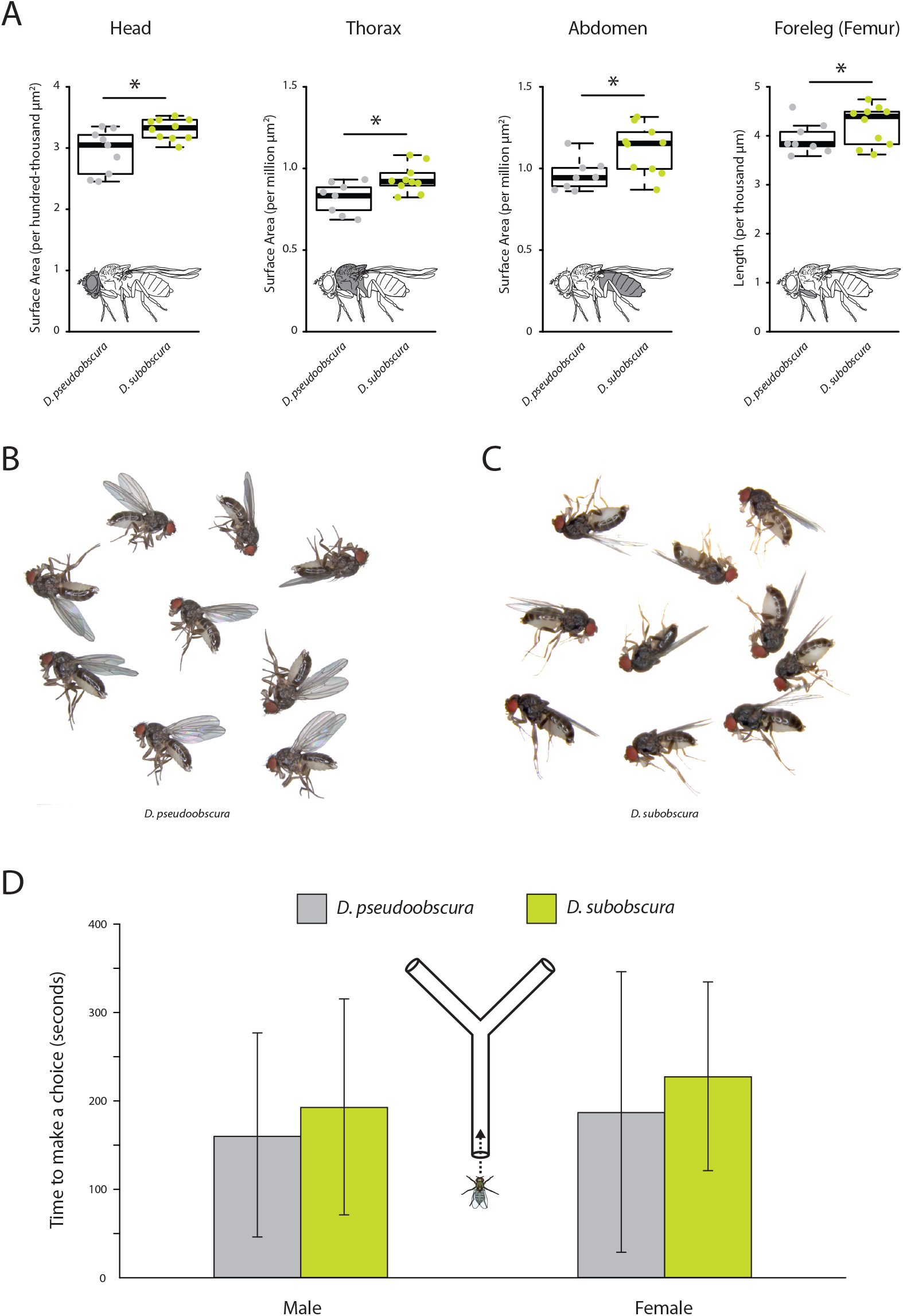
External morphometrics of two *obscura* species. Images of whole flies were collected to provide measurements on overall size differences between these two *Drosophila* species. While *D. subobscura* was significantly larger, this does not appear to explain the observed variation in eye or antenna sizes between these two species. (A) Lateral views of each fly species were measured to provide surface area estimates for head, thorax, abdomen and foreleg (femur) measurements. In all cases, *D. subobscura* females were significantly larger than *D. pseudoobscura* adults. (B) Lateral views of *D. pseudoobscura* adult females. (C) Lateral views of *D. subobscura* females. (D) Time metrics are shown for flies selecting between darkened or lightened sides of the Y-tube during phototaxis experiments.

**Supplementary Figure 2.**
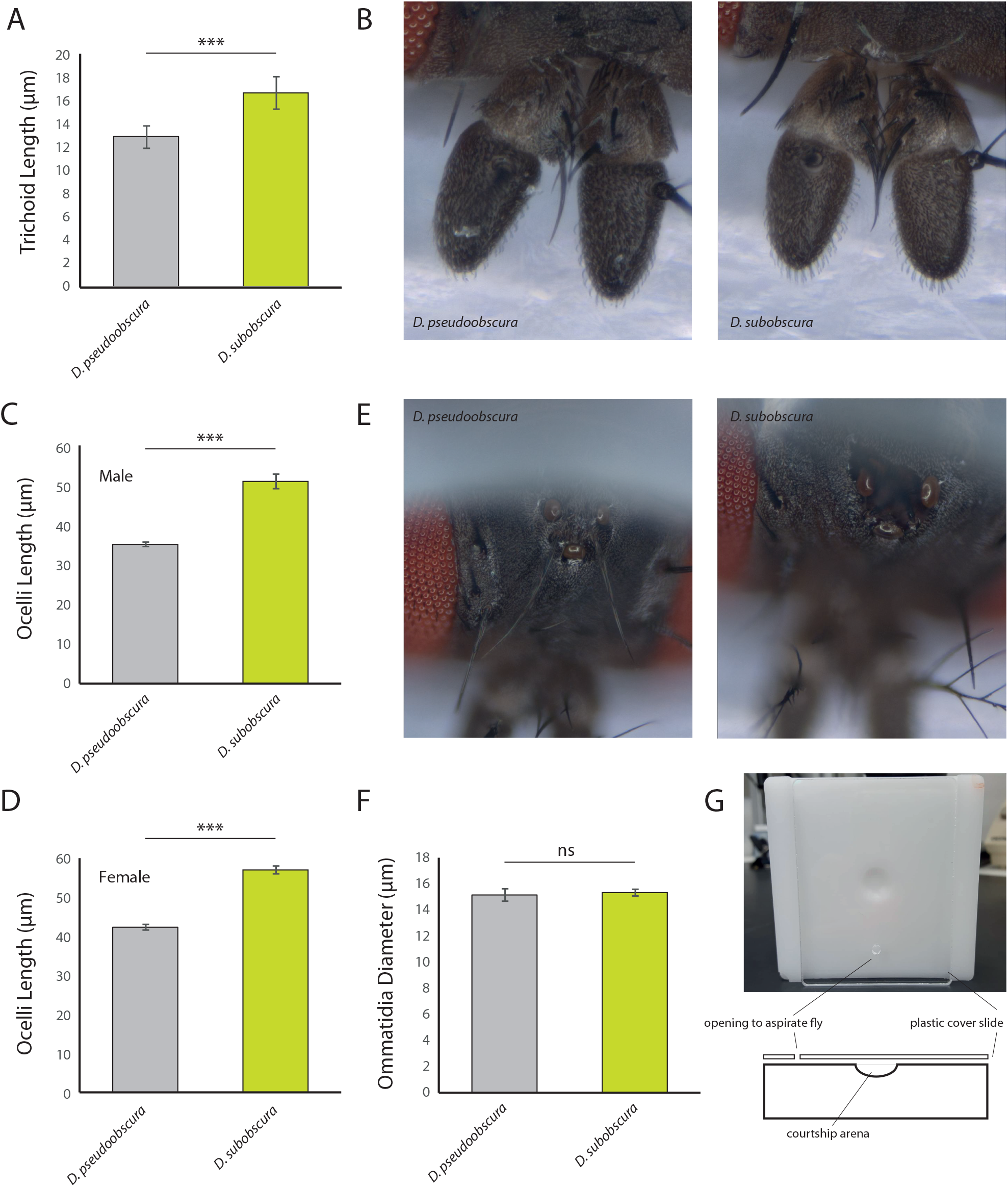
Additional metrics from sensory systems. (A) Here, we note a difference in trichoid length between males of the two *obscura* species, with *D. subobscura* having significantly longer sensillum than *D. pseudoobscura*. (B) Stacked focal images of the antenna from each species. (C-D) Average ocelli length from males and females of each species, where *D. subobscura* consistently had larger ocelli. (E) Stacked focal images of the ocelli from each species. (F) Average ommatidium diameters, where there was no significant difference between our two *obscura* species. (G) Courtship arenas used in this study.

## Notes

#### Summary of Updates

Supplementary Figure 2G updated, as well as the corresponding text (methods section: Analyses of courtship and mating behavior).

